# Pharmacological profiling of brain activity in zebrafish

**DOI:** 10.64898/2026.05.27.728255

**Authors:** Richard Kanyo, Ethan Smith, W. Ted Allison, Harley T. Kurata

**Affiliations:** Dept. of Pharmacology, Alberta Diabetes Institute, University of Alberta, 9-70 Medical Sciences Building, Edmonton, AB, T6G 2H7, Canada; Dept. of Biological Sciences, Centre for Prions and Protein Folding Disease, University of Alberta, Z 10-32 Biological Sciences Building, Edmonton, AB, T6G 2E9, Canada

**Keywords:** Kv7, KCNQ, potassium channel, CaMPARI, LOPAC1280, drug screen, zebrafish

## Abstract

**Background and Purpose:** Epilepsy is a neurological condition characterized by recurring seizures and neuronal hyperexcitability. Cell-based high-throughput screening applications have been essential for drug development and discovering novel biological processes. However, cell-based screens do not provide information on how drug-targeted pathways are integrated into a whole animal. Our objective was to develop and evaluate a screening application using zebrafish larvae to identify signalling mechanisms that modulate neural activity.

**Experimental Approach:** We developed an *in vivo* automated high-content screening assay using zebrafish larvae expressing the calcium sensor CaMPARI (calcium-modulated photoactivatable ratiometric integrator) in neurons. This assay can quantify neural activity of multiple individual larvae per well in a 96-well format. We quantified neural activity in 8725 individual larvae, in response to 1292 different drugs to identify molecules that protect against convulsant-induced neuronal hyperexcitability.

**Key Results:** The assay was effective at identifying drugs that target diverse neurotransmitter signalling systems. While some commonly used anti-convulsants (e.g. phenytoin, carbamazepine, valproic acid) had poor activity in the assay, Kv7 potassium channel activators were consistently effective (ICA-069673, ICA-27243, ICA-110381, retigabine, and ML213). Many compounds approved for treatment of other conditions, including amitriptyline (depression), cyclobenzaprine (muscle spasm), clomipramine (obsessive-compulsive disorder) and ganaxolone (seizures), also strongly suppressed excitability in the assay.

**Conclusion and Implications:** Neuronal CaMPARI expression in zebrafish larvae is a powerful tool for plate-based compound library screening to identify drugs that suppress hyperexcitability *in vivo*.

**Bullet Point Summary:** *What is already known:*

- CaMPARI is an integrative Ca^2+^ sensor that can be used to identify active neurons.
- Kv7 activators (retigabine, ML213, and ICA-069673) are effective at reducing convulsant-induced (4-AP) neuronal hyperexcitability.

*What this study adds:*

- An automated *in vivo* high-content drug screening assay to quantify neural activity.
- A series of drug targets that influence convulsant-induced hyperexcitability.

*Clinical significance:*

- Our new tool will help identify novel compounds and signalling mechanisms that could be pursued as therapeutic targets for diseases involving electrical hyperexcitability.

## INTRODUCTION

High-throughput screening of large compound libraries has led to the discovery of small molecules with therapeutic potential and tools to address fundamental biological questions. Various drugs to treat serious conditions such as cancer, epilepsy, or pain originated from screening large compound libraries using cell-based strategies, followed by optimization (Coussens et al., 2017; Kehne et al., 2017; Wu and Lu, 2026). However, a major drawback of *in vitro* cell-based screening is that findings do not inform on the integration of the targeted pathway in the whole animal. Lacking this physiological context likely contributes to the failures of many drugs at later stages of development, often because of a lack of target specificity, toxicity, or undesirable pharmacokinetics. In contrast, whole-animal models situate target receptors in differentiated cell types and native physiological settings, thereby generating a more complex view of how signalling pathways are integrated into general physiological function. However, mammalian models are time-consuming, expensive, and thus limit screening throughputs. The zebrafish larvae is an increasingly useful model that combines many benefits of cell-based and whole-animal assays. The zebrafish has 70% of its genome evolutionarily conserved with humans, and larvae are frequently used in translational research and to study fundamental biological processes (Rihel et al., 2010; Howe et al., 2013; Patton et al., 2021; Weinschutz Mendes et al., 2023; Zhao et al., 2023; Özcan and Rihel, 2026). The zebrafish larva has been especially useful as a model for epilepsy, owing to well-characterized seizure behaviours together with advances in calcium sensors for quantifying neural activity (Baraban et al., 2005, 2007; Akerboom et al., 2012; Fosque et al., 2015; Kanyo et al., 2020; Locubiche et al., 2024).

Epilepsy is among the most prevalent neurological disorders, affecting 1% of the global population. Roughly 30% of the patients exhibit drug-resistant seizures, due in part to the heterogeneity of underlying seizure mechanisms (Perucca et al., 2023). Therefore, efforts are ongoing to identify novel drugs and targets, and thereby broaden the number of available therapeutic alternatives. Common treatments target voltage-gated Na+ (Nav) channels and GABA-mediated neurotransmission, either by sensitizing GABA-A receptors (e.g., benzodiazepines or barbiturates) or by increasing GABA release (Sills and Rogawski, 2020). Kv7 voltage-gated potassium channels are more recently pursued therapeutic targets, recently highlighted by development of effective alternatives to retigabine/ezogabine such as XEN1101/Azetukalner (Blackburn-Munro et al., 2005; Orhan et al., 2012; French et al., 2023). While existing drugs continue to be optimized, there is a strong demand for the discovery of novel targets to influence neuronal firing. Therefore, methods to accelerate *in vivo* screening of drug actions on neural activity would be highly beneficial.

Zebrafish larvae enable compound library screening and incorporate important aspects of *in vivo* models, such as the blood brain barrier and drug metabolism (Rennekamp and Peterson, 2015). Locomotor activity as a surrogate measure of neural activity has been the most common approach for zebrafish larval screens, and has been used to study epilepsy, autism, and sleep (Baraban et al., 2007; Rihel et al., 2010; Hoffman et al., 2016; Patton et al., 2021). Genetically encoded calcium sensors such as GCaMP and GECO provide more direct quantification of neuronal firing (Mank and Griesbeck, 2008; Akerboom et al., 2012; Shaner et al., 2013; Zarowny et al., 2020). However, high-content drug screening using most calcium sensors in zebrafish is impractical because of animal movement and resource-intensive analysis. The calmodulin-modulated photoactivatable ratiometric integrator (CaMPARI) we have used in previous studies (Fosque et al., 2015; Kanyo et al., 2020, 2021, 2023), overcomes many of these challenges. CaMPARI photoswitches irreversibly from green to red fluorescence emission in the presence of high Ca^2+^ and 405 nm light, providing an integrated report on neural activity in freely moving organisms. The quantitative output is the ratio of red:green fluorescence emission intensities (Figure 1) and this ratiometric output somewhat normalizes against inter-individual differences in tissue orientation/depth and CaMPARI abundance.

**Figure 1.**
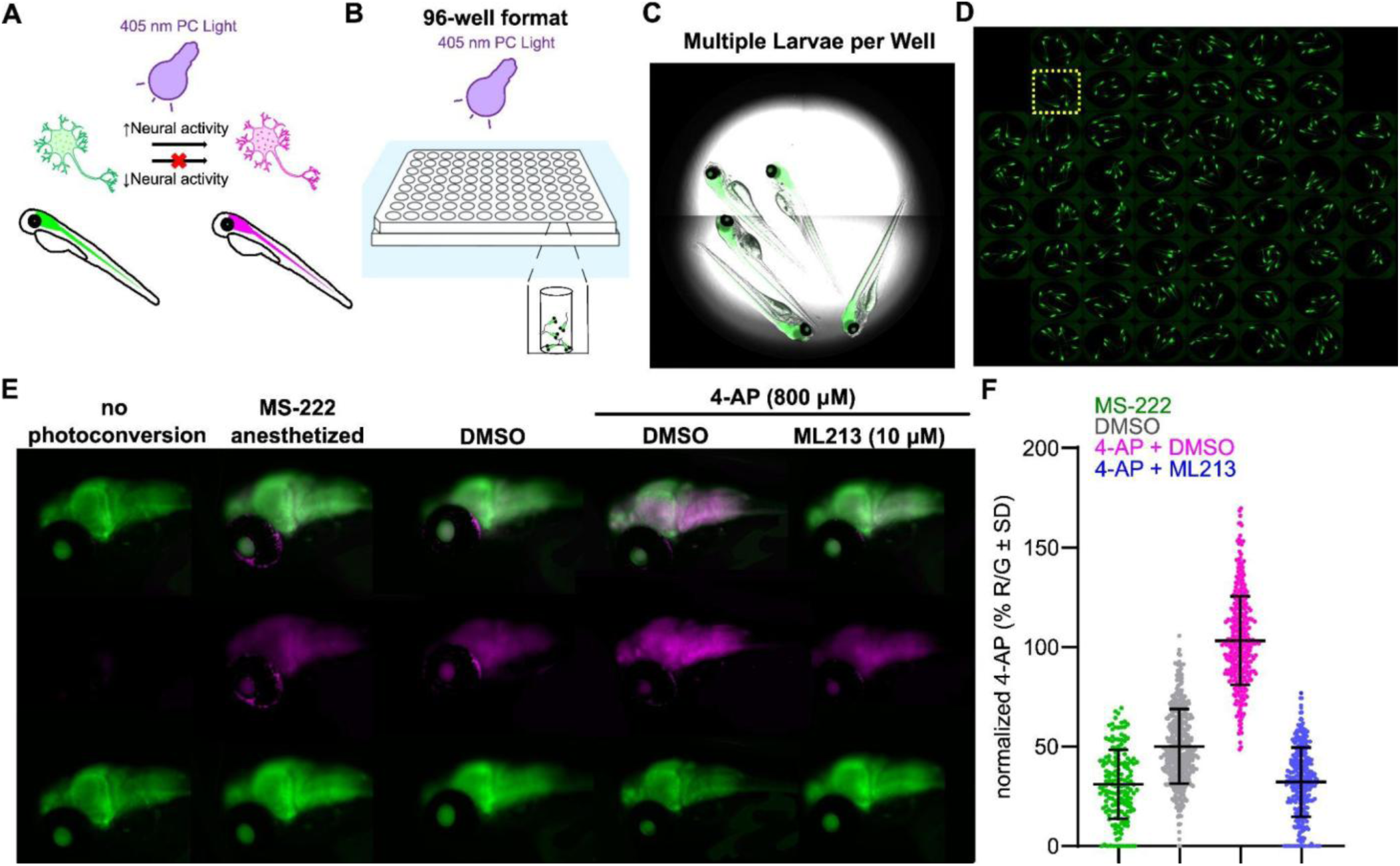
*In vivo* screen for modulators of convulsant-induced neural activity. Zebrafish larvae were engineered to express CaMPARI in CNS neurons. High-content imaging quantifies the impact of treatments on Ca^2+^ abundance as a proxy for neural activity. **A,** *Top*, CaMPARI photoconverts Ca^2+^ during exposure to photoconverting (PC) light. High Ca^2+^ together with PC light causes CaMPARI to switch from green to red fluorescence emission (pseudocoloured to magenta). *Bottom*, Increases in the red-to-green ratio (R/G) are interpreted to reflect neural activity. **B**, 96-well format automated high content microscopy enables imaging of several larvae per well. **C**, Representative bright-field image of a well with five anesthetized larvae (CaMPARI in the CNS). **D**, Image of a full 96-well plate illustrating CaMPARI fluorescence in the CNS of hundreds of larvae. The yellow square indicates one well for scale. **E**, Representative images of CNS CaMPARI emission (anterior to left) in indicated conditions: merged (*top*), magenta (*middle*), and green (*bottom*). **F**, Red:Green (R:G) ratio in individual larvae is normalized to the 4-AP response within the same plate. Error bars show standard deviation (SD). Mean R:G ratio, SD, and biological replicates (n) in the entire primary screen for the four control conditions are as follows: MS-222 (30.2 ± 16.8 %, n=170), DMSO (48.8 ± 18.0 %, n=447), 4-AP at 800 µM+ DMSO (100.0 ± 21.0 %, n=466) and 4-AP + ML213 at 10 µM (mean=31.6 ± 16.7 %, n=348). Panels A and B are modified sketches from (Kanyo et al., 2021).

In this study, we have adapted transgenic CaMPARI zebrafish as a medium-throughput *in vivo* drug screening tool for the identification of compounds with anti-convulsant/seizure properties. We used this tool to screen the LOPAC1280 compound library towards identifying drugs that influence chemical convulsant-induced changes in CaMPARI signal. Several compounds targeting neurotransmission were identified, and some are FDA-approved and used to treat a variety of conditions, including seizures, depression, and pain. Overall, our study establishes a useful combination of model organism and automated analysis to accelerate *in vivo* screening in early stages of drug development.

## MATERIALS AND METHODS

### Animal Ethics

Zebrafish were maintained in accordance with the University of Alberta’s Animal Care & Use Committee (Biosciences) and operated under the standards of the Canadian Council of Animal Care, as approved under AUP00000077.

### Drug preparation

ICA-27243, ICA-069673, ICA-110381, retigabine and ML213 were obtained from Tocris (Bristol, UK), and stock solutions were prepared in DMSO at 100 mM (ICA-27243, ICA-069673, ICA-110381 and retigabine) and 50 mM (ML213). 4-Aminopyridine (4-AP), carbamazepine, phenytoin, and valproic acid were obtained from Sigma-Aldrich. 4-AP (at 80 mM) and valproic acid (at 100 mM) were stored in water, while carbamazepine and phenytoin were stored in 50 mM DMSO stock solutions. Experimental solutions were prepared on each experimental day. The library of pharmacologically active compounds (LOPAC1280, Sigma-Aldrich) was obtained from the University of Alberta Advanced Cell Exploration core facility. Drugs were stored in sealed 396-well plates at 10 mM (DMSO) at -80 °C. For transport, compounds were aliquoted in 96-well plates and diluted to 200 μM in E3 zebrafish media to generate experimental/working plates for the primary screen (Figure S1).

### Drug screen via CaMPARI assay

Drug treatment of zebrafish larvae was set up in a 96-well plate format, similar to a previous study (Kanyo et al., 2021). The transgenic zebrafish line Tg(elavl3:CaMPARI [W391F + V398L])^ua3144^ was generated as previously described (Kanyo et al., 2020). Larvae were placed in the central 64 wells of a 96-round-well plate (Sigma-Aldrich) and acclimatized in E3 for at least one hour prior to treatment (Figure S1). All treatment groups contained a final concentration of 0.3% DMSO. Drugs from the LOPAC1280 were applied at 30 μM in a total of 150 μl, unless otherwise indicated. Each experimental/working plate contained 40 compounds from the LOPAC in combination with the convulsant 4-AP at 800 μM, and included the following control groups: DMSO (vehicle control), 4-AP at 800 μM plus DMSO, 4-AP plus ML213 at 10 μM, and a group that was not exposed to the photoconversion (PC) light (Figure 1S). Some experimental/working plates also included groups that were anesthetized with 920 μM MS-222. The treatment lasted 30 min, followed by photoconversion of CaMPARI. While Figure 1S describes the setup for the primary screen (Figure 2), available wells were used for treatments in other experiments.

**Figure 2.**
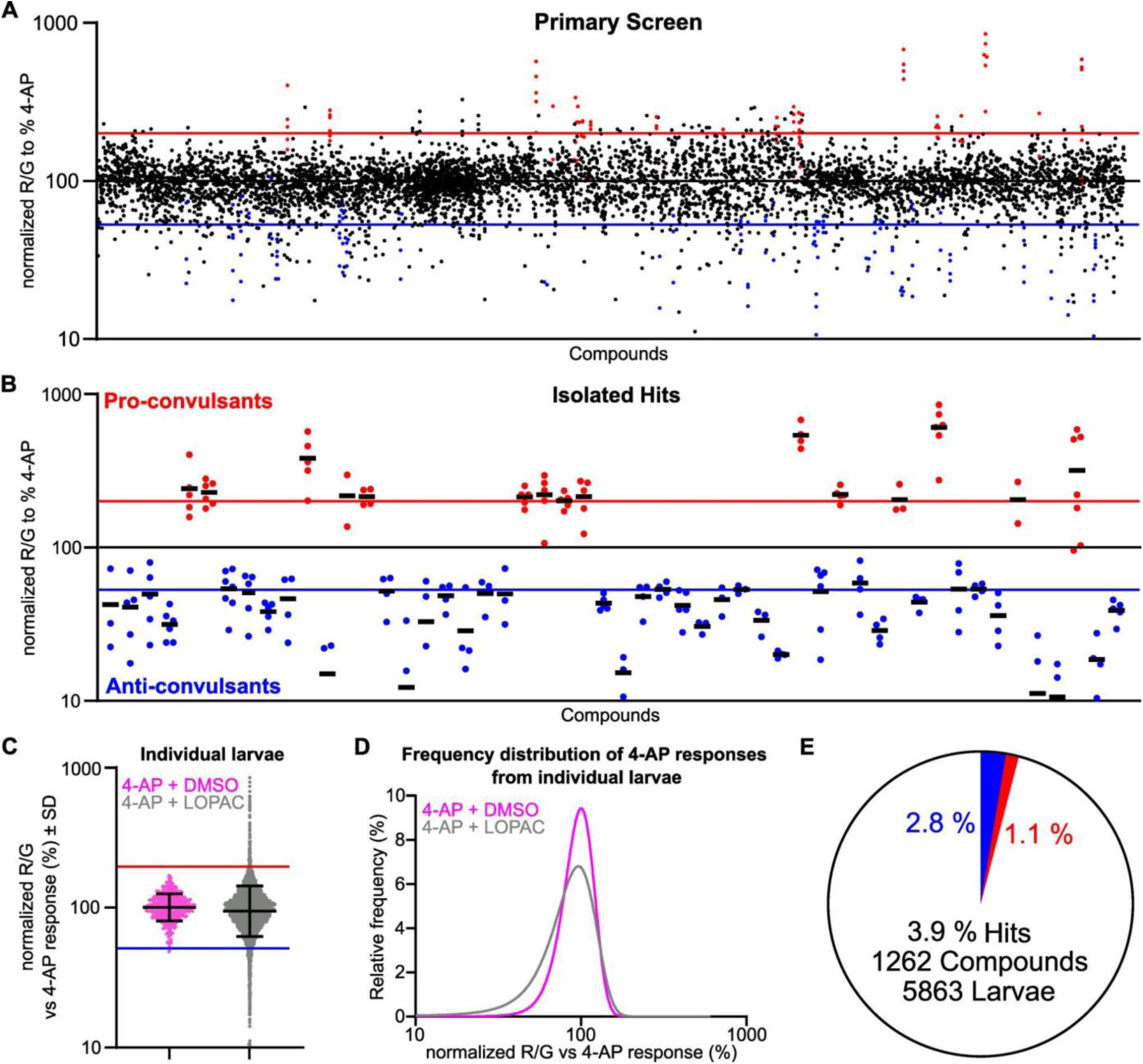
Primary screen for compounds that modulate convulsant-induced CNS activity in zebrafish. Zebrafish larvae at four days post-fertilization were treated for 30 min with 4-AP at 800 μM, along with a compound from the LOPAC1280 library (30 μM). The ratio of red (R) and green (G) fluorescence (R:G) is a proxy for neural activity. Each data point reflects one zebrafish larva brain. Data were normalized to the mean R:G ratio in 4-AP treatment in each working plate, and each plate tested 40 drugs from the library (see Figure S1). **A**, The normalized red:green ratio for every zebrafish brain in our primary screen. Results from all fish exposed to each compound are shown. Blue and red lines depict cut-offs for hits with anti- and pro-convulsant properties, respectively. **B**, Drugs from panel A that met the defined “hit thresholds” are highlighted. The threshold for pro-convulsant activity (amplifying 4-AP) was set at >200 % of the mean 4-AP response (3 SD above the mean of 4-AP alone). The threshold for anti-convulsant activity was set at 5 % less than the baseline R/G ratio in the DMSO vehicle (<54% of the 4-AP response; see Figure 1F). **C**, All data points from individual brains treated with 4-AP alone, or with a LOPAC compound, revealing the greater range of R:G ratio among larvae treated with test compounds (mean +/- SD). **D**, Frequency distribution (5 % bins) of all LOPAC-treated samples versus 4-AP (no test compound). **E**, Percentage of drugs identified as anti-convulsant (blue), or pro-convulsants (red).

Photoconversion leads to irreversible photoswitching of CaMPARI from green to red fluorescence emission (Figure 1A). Following the indicated treatments, larvae were exposed to the photoconversion light for 20 s (Figure 1B), at a distance of 2 cm from the 405 nm LED array (Loctite), as previously described (Kanyo et al., 2020). Next, larvae in the 96-well plate were washed with E3 media and 0.24 mg/mL tricaine methanesulfonate (MS-222, Sigma) to immobilize the animals for imaging.

Image acquisition of 96-well plates was done using an ImageXpress high-content microscope with a 4.0x objective (Molecular Devices, LLC, US), generating 4 images per well that were stitched together. Three imaging channels were used: FITC channel to visualize the green fluorescing non-photoconverted CaMPARI, Cy3 for the red fluorescing photoconverted CaMPARI, and DAPI for masking the yolk. To isolate the zebrafish brain as a segment in the images, a mask was created using MetaXpress 6 software provided by the ImageXpress high-content imager. Briefly, a protocol was established to identify regions of above-background emission intensity in the FITC channel that are similar in size to the brain area. Autofluorescence of the yolk sac can interfere with the detection and quantification of CaMPARI expressed in the nervous system. Autofluorescence from the yolk sac also appeared in the DAPI channel. Thus, the DAPI channel was used to create a mask by setting a fluorescence threshold to clearly detect the yolk of each larva, and then this area was subtracted from the FITC and Cy3 channels, yielding the mean fluorescence intensities of FITC and Cy3 of the zebrafish brain area. This approach was used to independently image and quantify multiple zebrafish brains in each well. Background fluorescence in the Cy3 channel was subtracted using larvae that were not treated with the PC light, which were included as controls on every plate (Figure S1). The ratio of red:green fluorescence intensity was quantified and interpreted as relative neural activity, as previously defined (Fosque et al., 2015). For our study, the red:green ratio is normalized as a percentage of the mean value of all the larvae treated with 4-AP on each experimental 96-well plate.

### Data analysis

Guidelines for this study are aligned with the British Journal of Pharmacology and recommendations previously established (Curtis et al., 2018). While our group sizes were comparable to the test compounds, the control groups typically consisted of more replicates to increase the reliability of our findings across thousands of samples across our drug screen. For efficiency and resource limitation, treatment groups of experiments outside of our primary screen were included and compared to the controls in the same working plate (Figure S1). The groups indicated across plates were compiled to strengthen our conclusions and to cross-compare individual plates. Note that the group size was also affected by the variable success of the drug screen due to fish breeding, drug removal, and imaging individual wells. Randomization was not applied because the drugs were labelled with numbers and the output was automated. This ensured that the experimentalist was blinded to most treatments. Each individual larva was treated as a biological replicate reported as “n”, or group size. Data is presented as mean ± standard deviation (SD) or standard error of margin (SEM), as indicated. Statistical analysis was performed with independent replicates, not technical replicates, with each group size at least n = 5. If only three or four replicates were analysed it was due to limitation of the compound available to us and the loss of some samples, which we recognize is a limitation of our study. P values < 0.05 were considered significant for all statistical tests. The Shapiro-Wilk test was used to assess the normality of all data used for statistical tests. Student’s t-test was used in cases for comparison of two groups. In multiple-group comparisons with parametric variables, an F-test was performed first, and post hoc tests were conducted if the F-test p-value was <0.05 and there was no significant heterogeneity of variance. One-way ANOVA followed by Dunnett’s multiple-comparison test was used for multiple-group comparisons against a single reference. For Figure 6B, a two-way ANOVA followed by a post-hoc Dunnett’s test was used. On our primary screen, larvae may be positioned poorly, hindering proper recognition by our software and reliable measurements. Therefore, for our primary screen in Figures 1 and 2, we objectively excluded outliers using Grubbs’ test at α=0.2.

**Table 1.** Drugs identified as anti-convulsants. Shows all confirmed hits on the secondary screen that are used to treat indicated medical conditions, which are anti-convulsants.

| Compound Name | Medical Treatment |
| --- | --- |
| Amitriptyline | major depression, neuropathic pain, migraines, insomnia, adhd, psychotic disorders, anxiety, seizures |
| Maprotiline | major depression, panic disorder, neuropathic pain |
| Ivermectin | anti-parasite medicine |
| Oxybutynin | overactive bladder |
| Trifluoperidol | schizophrenia |
| Propionylpromazine | anti-psychotic |
| Cyclobenzaprine | muscle spasm |
| Ganaxolone | anti-seizure medicine |
| Riluzole | amyotrophic lateral sclerosis (ALS) |
| Clomipramine | obsessive-compulsive disorder (OCD) |
| Imipramine | major depressive disorder, nocturnal enuresis, neuropathic pain, panic disorder |
| Trimipramine | depression |
| Tetracaine | local anesthesia, seizure |
| Vancomycin | antibiotics |

**Table 2.**
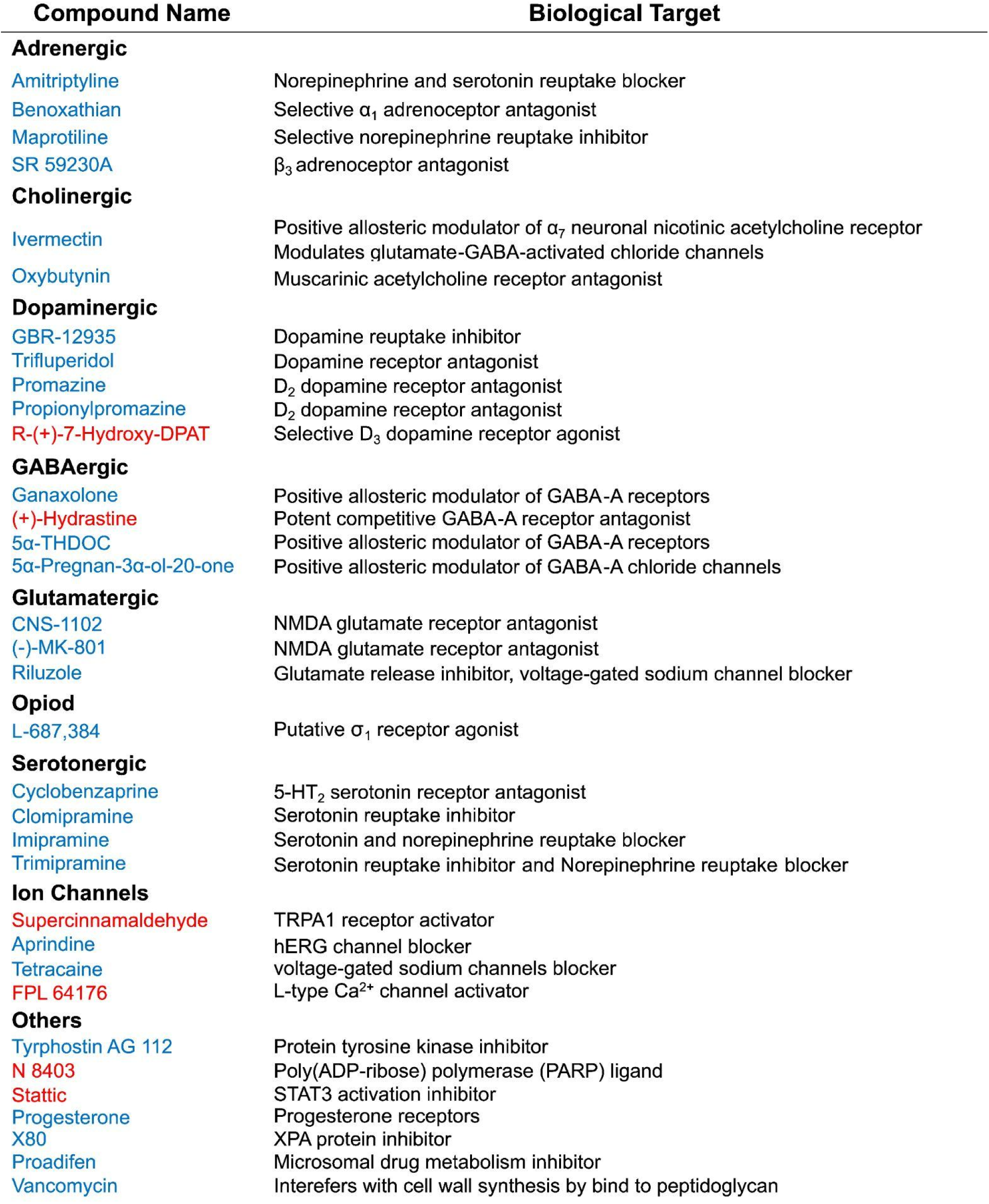
Anti- and pro-convulsants target diverse signaling mechanisms. Shows all confirmed hits on the secondary screen, organized into groups of compounds by biological targets. Anti-convulsants are in blue; pro-convulsants are in red.

## RESULTS

Screening approaches in zebrafish have typically used locomotion-based assays as a proxy of seizure-like behaviour. We sought to develop an automated screening assay to directly quantify neural activity using a genetically encoded calcium sensor (Mank and Griesbeck, 2008; Akerboom et al., 2012; Zarowny et al., 2020). We developed approaches for plate-based high-content imaging of transgenic CaMPARI-expressing zebrafish to enable screening of small-molecule compound libraries. We used this ‘CaMPARI assay’ to screen for anti-seizure compounds, and we tested it against traditional anti-seizure drugs and a series of emerging compounds targeting voltage-gated Kv7 potassium channels, which have been found to have anti-convulsant properties.

### Compound library screening using the CaMPARI assay

We executed a primary screen to determine the effects of compounds in the LOPAC1280 library (Figures 1 and 2). Zebrafish larvae expressing CaMPARI in the CNS were treated with convulsant 4-AP and various test compounds in clear-bottom 96-well plates, followed by photoconversion, anesthesia, and high content imaging in the same plates (Figures 1B,S1). Each well contained multiple larvae (Figure 1C), yielding hundreds of individual larvae for separate quantification per plate (Figure 1C,D,S1). Each plate was set up with two to four wells of each of the various control conditions. Three or four wells of DMSO (vehicle) treated larvae provided a baseline measure of CaMPARI signal from vehicle control animals. Three or four wells were treated with 4-AP alone with no test compound, as a negative control to describe the change in CaMPARI signal in response to the chemical convulsant drug. Finally, as a positive control for anti-convulsant drug actions, between two and four wells were treated with 4-AP along with 10 μM ML213, an established Kv7 activator compound that powerfully suppresses the extent of CaMPARI photoconversion, indicating strong suppression of neural excitability (Kanyo et al., 2020). A typical plate layout with various treatment conditions is shown in Figure S1. Except during rare exceptions when larvae died in the well, each experimental well had 3 to 7 larvae. Treatment with 4-AP (800 μM) for 30 min induced a roughly 2-fold increase in CaMPARI photoconversion relative to the DMSO control (Figure 1E,F), indicating increased neural excitability. This is apparent in exemplar images in Figure 1E, as the red emission component of CaMPARI increases in intensity in the 4-AP condition, indicating a larger amount of the photoconverted CaMPARI, and thus higher intracellular calcium levels in neurons. The combined treatment of 4-AP and ML213 reduced neural activity to ∼35%; this was a positive control used on all plates, and demonstrates that certain drugs can prevent convulsant (4-AP)-induced hyperexcitability. The most important development in this assay that enabled screening of large numbers of drugs and larvae was the automation of imaging and analysis. Previous studies used manual quantification of CaMPARI signals from confocal microscopy images (Kanyo et al., 2020). To markedly improve throughput in a plate-based format, we adapted the assay to use basic fluorescence microscopy with a high content plate reader. Essential innovations for this included automated detection and masking of autofluorescence from the yolk sac, paired with automated detection and quantification of fluorescence from individual larvae in each well (see Methods).

To screen the LOPAC library, we treated zebrafish larvae with a standard concentration (30 μM) of each drug, together with 4-AP (800 μM), for 30 min followed by photoconversion, anesthesia, and imaging. In our primary screen, we assessed neural activity in the brains of 8725 individual zebrafish larvae, in response to 1280 different drugs (Figure 2A,B). For anti-convulsant activity, we arbitrarily defined a “hit” as treatments leading to a R:G (red:green) CaMPARI ratio close to the corresponding DMSO control (no 4-AP), indicating reversal of 4-AP-induced hyperexcitability (Figure 1F; <54 % of the 4-AP-induced response). In addition, multiple compounds exhibited pro-convulsant effects, and we defined a threshold of 3 standard deviations above the mean 4-AP-induced response. The data points of R:G ratio in all zebrafish controls followed a Gaussian distribution with the distribution of R:G ratio in LOPAC-treated fish exhibiting a broader distribution, reflecting the effects of pro- and anti-convulsant drugs (Figure 2C,D). The frequency distribution also shows that compounds in the LOPAC library with anti-convulsant properties appeared more frequently than pro-convulsant compounds (Figure 2D). Based on these thresholds, 3.9% of drugs in the library were classified as hits and considered for further validation. Among these, 2.8 % exhibited anti-convulsant activity (reduced 4-AP-induced effects), while 1.1 % exhibited pro-convulsant activity (increased neural activity above 200 % of the 4-AP level) (Figure 2E). Eighteen compounds could not be quantified because they were toxic to larvae or were autofluorescent in the spectral range used for imaging. Overall, these experiments identified 49 hits for further investigation/validation in a secondary screen.

### Activity of Kv7-targeted activator compounds in the CaMPARI screen

Many recent efforts have targeted the development of Kv7 potassium channel activators as anti-seizure medications, although these drugs are poorly represented in the LOPAC library (Miceli et al., 2018; Perucca and Taglialatela, 2025). Therefore, we also took advantage of the faster throughput of the plate-based CaMPARI assay to determine the concentration response of a variety of experimental Kv7 activator drugs (Figure 3). Kv7 activators (eg. retigabine, ML213, and ICA-069673) were previously characterized with manual approaches and would serve as an additional control to validate our screening tool (Kanyo et al., 2020). All Kv7 activators reduced the R:G ratio, in the presence and absence of 4-AP (Figure 3A). Activators targeting the Kv7 channel pore (i.e. retigabine and ML213), were the most potent and effective in reducing 4-AP-induced neural activity (Figure 3B). In contrast, potentiators targeting the Kv7 voltage-sensing domain (VSD; i.e. ICA-27243, ICA-069673 and ICA-110381) required higher concentrations of 10 μM and 30 μM to achieve a clear reduction in the 4-AP stimulated R:G ratio (Figure 3B). Differences between pore- and VSD-targeted Kv7 drugs are also apparent in the absence of 4-AP. While 10 μM retigabine or ML213 clearly reduced the R:G ratio, VSD-targeted drugs were less effective. Together, these findings demonstrate that CaMPARI screening is sensitive to a range of Kv7 activator drugs in addition to established drug classes that are better represented in the LOPAC library.

**Figure 3.**
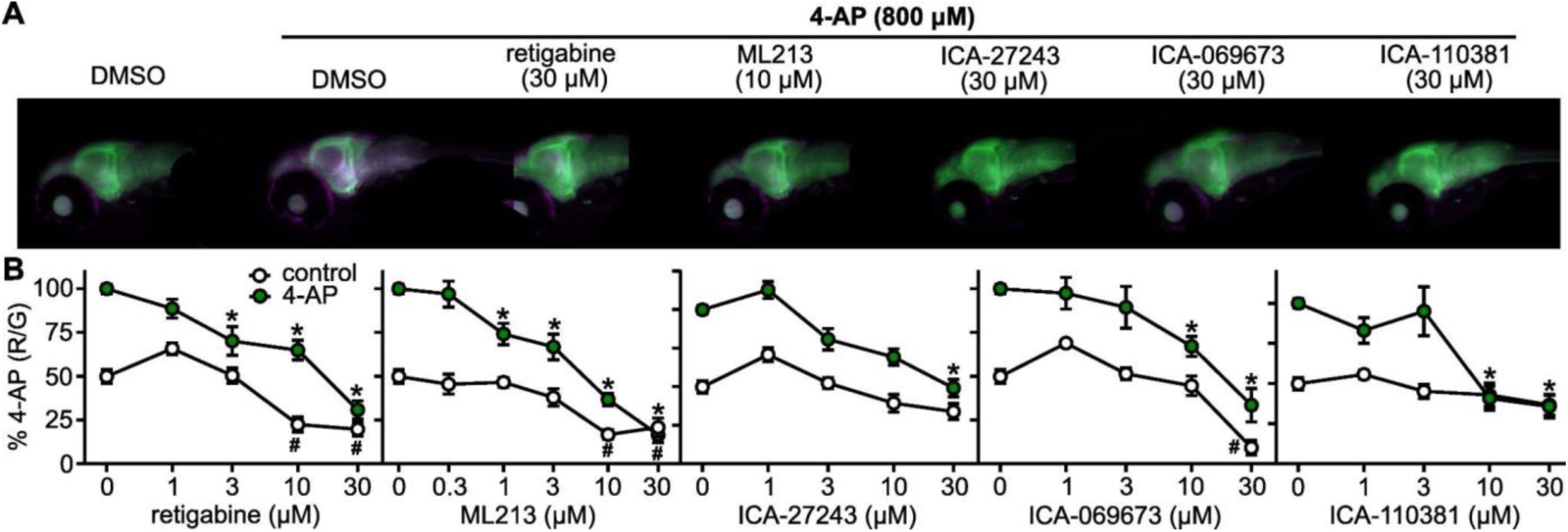
CaMPARI assay to validate diverse Kv7/KCNQ-targeted activators. Previously identified Kv7/KCNQ activators were tested with the CaMPARI assay for *in vivo* validation with zebrafish larvae. **A**, Representative images merged channels of red (pseudocoloured magenta, Cy3) and green (FITC) CNS localized CaMPARI signal, under the indicated conditions. **B**, Dose-response relationships in the presence (green) and absence (white) of 4-AP. The red:green ratios are normalized to the 4-AP + DMSO condition. * indicates statistical significance vs 4-AP. # indicates statistical significance at p<0.05 vs control. (ANOVA, followed by Dunnett’s post hoc test). Biological replicates are at n=5-35.

### Activity of established anti-convulsants in the CaMPARI screen

It is also important to consider established drug classes for which the assay is not particularly well-suited. For example, despite widespread use for seizure treatment in humans, several anti-seizure drugs in the LOPAC library, such as carbamazepine, phenytoin, and valproic acid did not meet activity thresholds in the zebrafish assay. One consideration is that 30 μM was the maximum concentration used in the plate-based assay, which may have been insufficient for some of these drugs to elicit a strong response. For comparison, carbamazepine, phenytoin, and valproic acid are reported to have IC50s against various Nav channels in the range of >100 μM, ∼10 μM, and >500 μM, respectively. Accordingly, we tested several higher concentrations of each drug in the presence of 4-AP (Figure 4). In general, these drugs had modest or absent effects, up to concentrations of 1 mM, regardless of whether they were co-applied with 4-AP (Figure 4A). This suggests that although the dose-response overlapped with the reported range of therapeutic concentrations (Wu and Lim, 2013; Acikgoz et al., 2016; Tseng et al., 2020), the CaMPARI assay using zebrafish may not be ideal to test Nav channel blockers. Another consideration may be that the assay involves acute drug treatments (∼30 minutes), which may not capture the full mechanism of action of certain anti-seizure medications.

**Figure 4.**
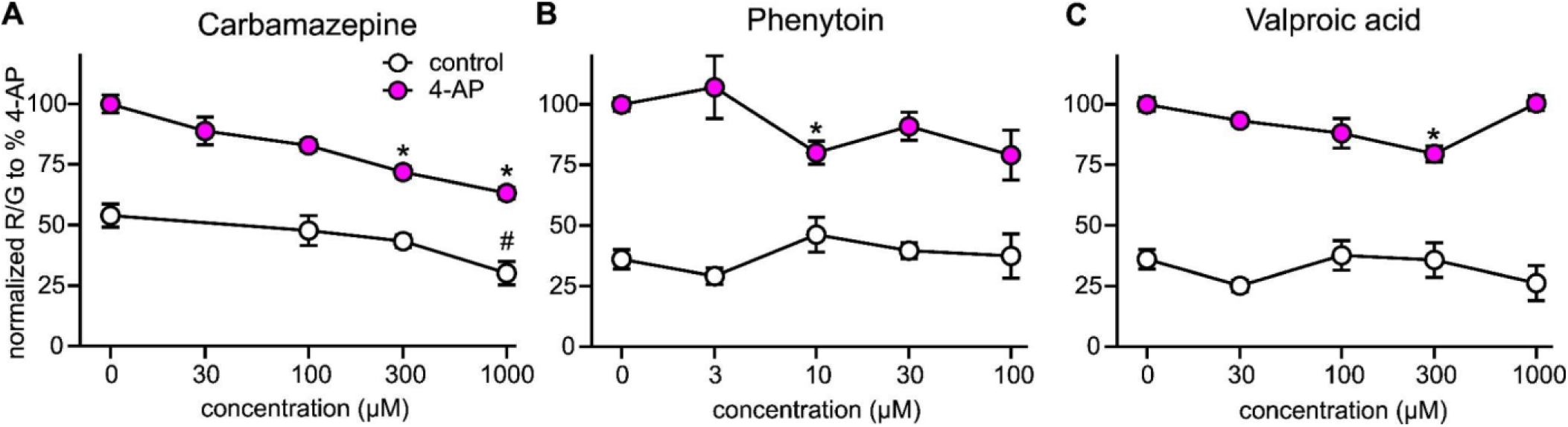
The CaMPARI assay exhibits modest responses to certain established anti-seizure drugs. Zebrafish larvae at four days post-fertilization were treated with carbamazepine (**A**), phenytoin (**B**) or valproic acid (**C**) in the presence (magenta) or absence of 4-AP (white) at 800 μM for 30 min. The red:green ratio is normalized to the 4-AP plus DMSO condition. *, marks statistical significance vs 4-AP. #, indicates statistical significance at p<0.05 vs control (ANOVA, followed by Dunnett’s post hoc test). Biological replicates are n=5-37.

### Confirmation of anti- and pro-convulsant compounds in the CaMPARI screen

The primary screen in Figure 2 typically included 3-7 larvae per drug condition, and we sought to validate findings with independent measurements in a secondary screen using identical conditions. We compared the secondary screen and primary screen findings to assess the quality of the CaMPARI assay (Figure 5). The secondary screen focused only on hits identified during the primary screen and tested each compound for statistically significant effects relative to the 4-AP convulsant condition of each plate. The overall quality of the assay was assessed by the correlation between the mean values from the primary and secondary screens, including compounds that were not confirmed, which yielded a correlation coefficient of *R* = 0.77 (Figure 5A). Among the confirmed anti- and pro-convulsants, the mean R:G ratio values were similar between the primary and secondary screens (Figures 5B,C, S2). This approach confirmed 67% of total hits from the primary screen (Figure 5D). Among the anti-convulsant hits, 80% of drugs were confirmed in the secondary screen (Figure 5E). Together, these results indicate that the assay is a robust screening tool for automated *in vivo* quantification of neural activity, and is especially reproducible for the identification of anti-convulsant drugs. 33 compounds were confirmed to affect 4-AP-induced neural activity, and 28 of those had anti-convulsant properties (Figure 6). Interestingly, we identified several FDA-approved compounds used for various treatments in addition to seizures (Figure 6A, Table 1). Drugs confirmed in the secondary screen were organized by drug class and target (Figures 6B,C,S4, and Table 2). These anti-convulsants target neurotransmission with diverse targets, including adrenergic, cholinergic, dopaminergic, serotonergic, GABAergic, and glutamatergic signalling cascades. We benchmarked the effects of anti-convulsants to the established Kv7 activator ML213, previously shown to be highly effective at suppressing 4-AP-induced hyperexcitability (Figure 1). At 30 μM, three compounds (GBR-12935, riluzole, and clomipramine) reduced the R:G ratio further than ML213.

**Figure 5.**
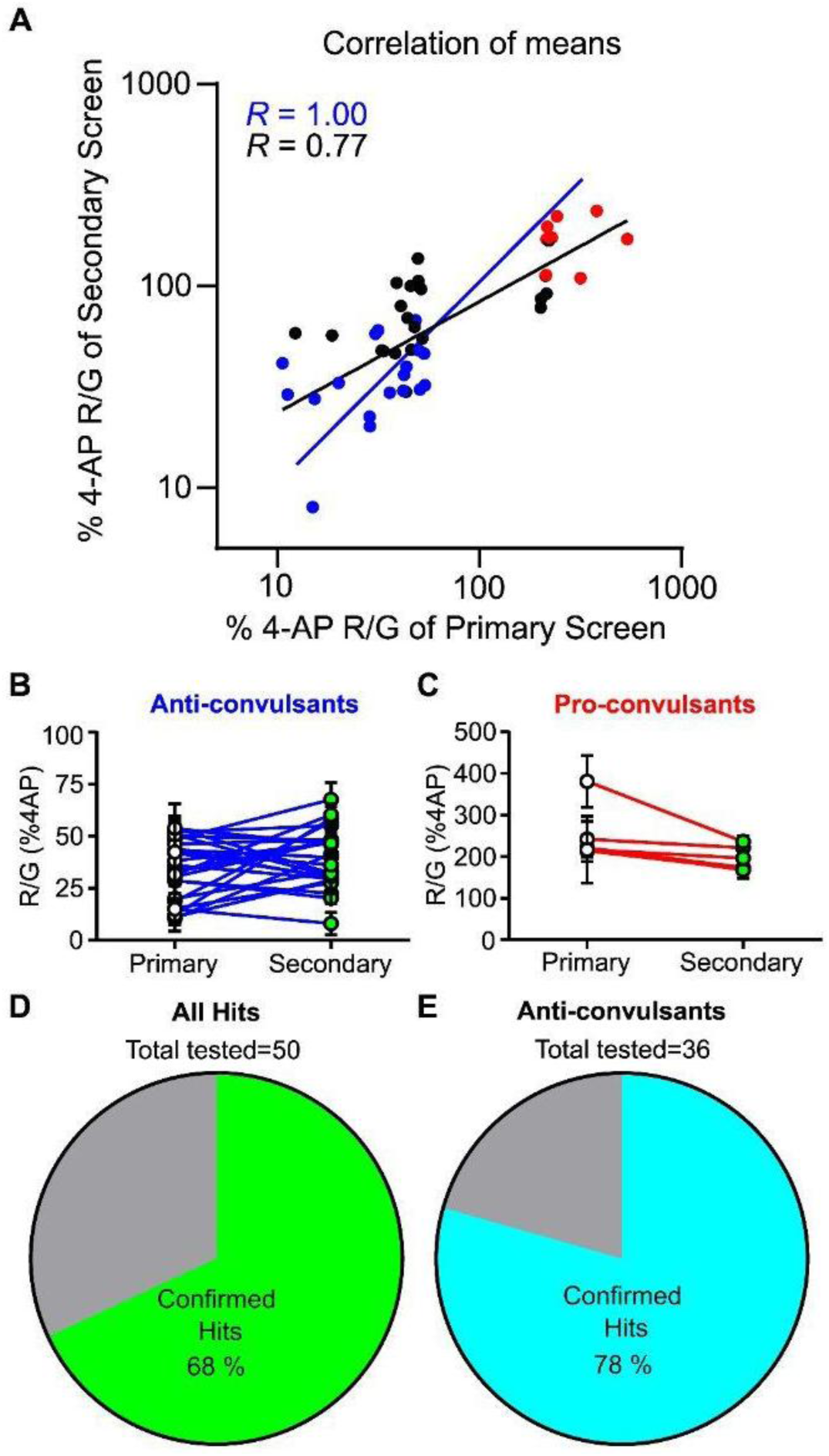
The secondary screen reveals reproducibility of the CaMPARI assay. A secondary confirmatory screen with identical treatments as the primary screen was executed for primary screen hits. Zebrafish larvae (4 days post-fertilization) were treated with LOPAC1280 test compounds (30 μM) for 30 min, with 4-AP (800 μM). **A**, Assay quality was assessed by the correlation between the means of all hits (confirmed and non-confirmed) from the primary and secondary screens. The red-to-green ratio was normalized to the 4-AP response on the same working plate. Confirmed anti-convulsants are blue, confirmed pro-convulsants and black dots are non-confirmed hits. **B** and **C**, Comparison of R:G ratios in all confirmed anti-convulsants (blue line; **B**) and pro-convulsants (red line; **C**) in primary and secondary screens. Error bars indicate ± SEM. Statistical significance at P < 0.05 was used to confirm hits in the secondary screen (ANOVA followed by Dunnett’s post hoc test, for each drug treatment compared with 4-AP alone, n=5-23).**D**, Percentage of hits confirmed by the secondary screen. **E**, Percentage of hits with anti-convulsant properties confirmed by the secondary screen.

In the secondary screen, we also assessed drug responses in the absence of 4-AP, to determine drug effects in freely swimming animals without a seizure insult (Figure 6B,C). In the absence of 4-AP, most drugs had negligible effects on the R:G ratio, although 6 of 28 drugs clearly reduced the R:G ratio (ivermectin, GBR-12935, riluzole, and all GABAergic positive allosteric modulators confirmed in our secondary screen: ganaxolone, 5α-THDOC, and 5α-Pregnan-3α-ol-20-one) (Figures 6B,S3). 5α-THDOC was among the most potent anticonvulsants against 4-AP induced hyperexcitability, with effects apparent at a concentration 1 μM in the presence of 4-AP (Figure S4). Lastly, among the compounds with pro-convulsant properties ((+)-hydrastine, supercinnamaldehyde, FPL 64176, N 8403, and stattic) only supercinnamaldehyde and stattic did not alter the R:G ratio in the absence of 4-AP (Figure 6C). These findings demonstrate that the CaMPARI assay can reliably identify compounds with anti- and pro-convulsant properties and is sensitive to compounds targeting a diverse set of neurotransmitter signalling pathways.

**Figure 6:**
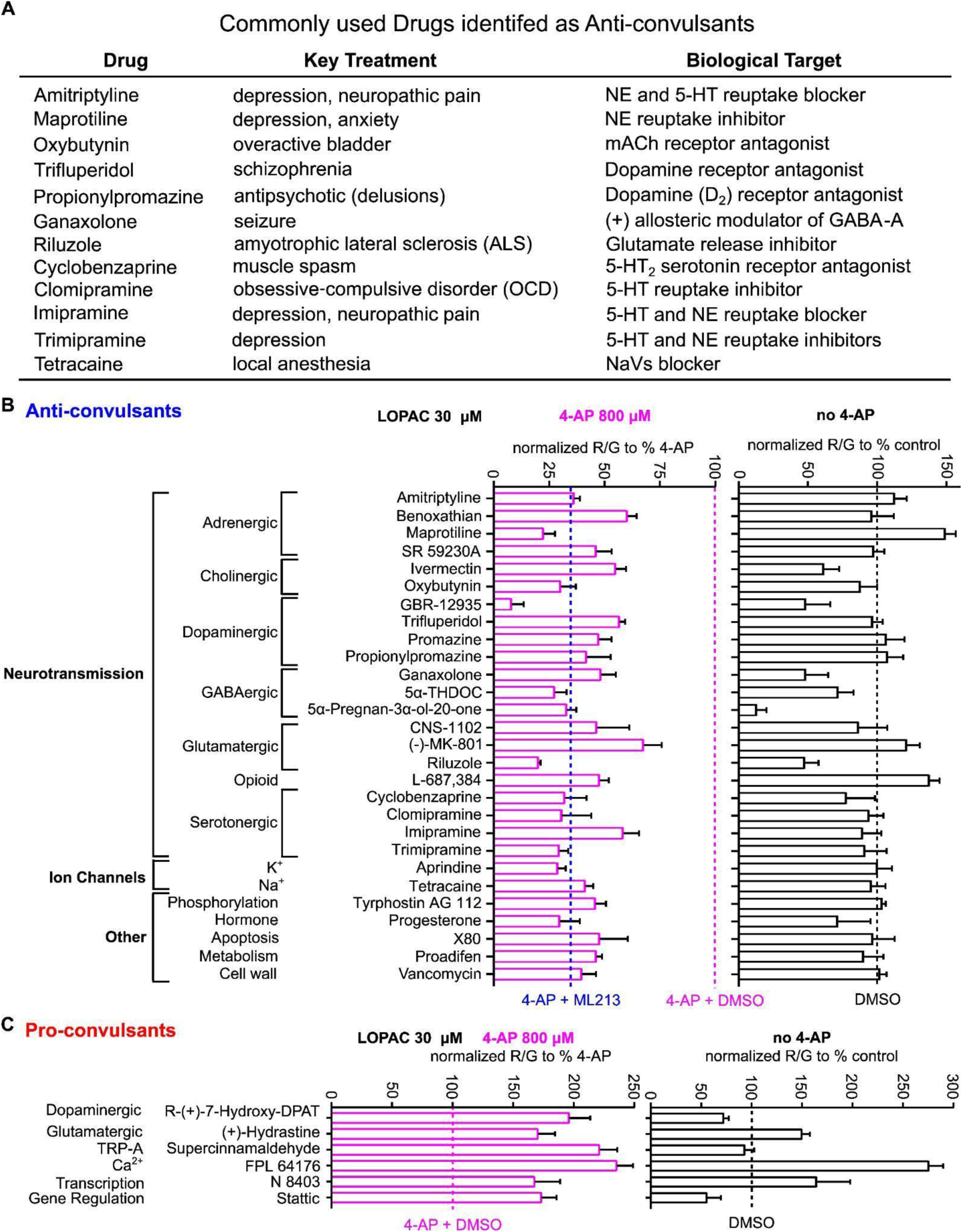
Independent secondary screen reveals diverse compounds with anti-, pro-convulsant properties and effects under basal conditions. Zebrafish larvae (4days post-fertilization) were treated with LOPAC1280 test compounds (30 μM) for 30 min in the presence (magenta) or absence (black) of 4-AP 800 μM. Confirmed hits are organized by compound class and drug target, as indicated and described in Tables 1 and 2. **A,** Hits from the LOPAC1280 that are FDA-approved. **B**, Classification of compounds with anti-convulsant properties. **C**, Classification of compounds with pro-convulsant properties.The R:G ratio is normalized to the mean 4-AP response (B,C; *left*) or the basal activity in DMSO alone (B,C; *right*) on the same working plate. Statistical significance at P < 0.05 was used to confirm hits (4-AP co-treated with LOPAC compounds were compared with 4-AP from the same working plate, one-way ANOVA followed by Dunnett’s post hoc test, n=5-23).

## DISCUSSION

High-throughput screening is an essential component of drug discovery and development. For instance, small molecules that modulate Kv7 channel activity have been identified in large compound libraries using a cell-based *in vitro* thallium influx or rubidium efflux assay followed by optimization in structure-activity relationship studies (Yu et al., 2010, 2011, 2016). While cell-based *in vitro* assays are advantageous in terms of economy and practicality, they do not account for complexities and opportunities that emerge in intact organisms. *In vivo* approaches are usually impractical and burdensome due to their complexity and limited throughput. Zebrafish larvae have been a useful animal model for implementing screening strategies that balance some of these pros and cons. Zebrafish larvae are especially popular for modeling epilepsy and for identifying seizure-resistant mutants to elucidate basic biological processes or physiologically relevant pathways for drug targeting (Baraban et al., 2007; Rihel et al., 2010; MacRae and Peterson, 2015). Here, we describe an automated *in vivo* drug-screening application that uses CaMPARI as an indicator to measure neural activity in the brains of individual zebrafish larvae. We used this innovation to screen over 8000 zebrafish brains and 1292 compounds. This will improve understanding of how neuronal excitability is integrated into the whole animal and inform therapeutic approaches targeting epilepsy.

Current treatments targeting seizures are prone to side effects or are ineffective in many patients (Kwan and Brodie, 2000), which creates a high demand for developing new drugs. Kv7 channels encode the neuronal M-current, and drugs targeting these channels have been developed relatively recently, including retigabine/ezogabine and XEN1101/azetukalner (Gunthorpe et al., 2012; French et al., 2023). To validate our *in vitro* findings on the mechanism of Kv7 modulators in a physiological context, we have previously used transgenic zebrafish larvae expressing CaMPARI (Kanyo et al., 2020, 2023). Instead of manual confocal microscopy, we optimized conditions for high-content imaging to set up our drug screen. We tested our previously published Kv7 potentiators while adding two more compounds in Figure 3 that are similar in structure. Briefly, aligned with our previous findings, the CaMPARI *in vivo* drug screening tool appears highly responsive to Kv7 potentiator drugs tested. In contrast, less-specific traditional antiseizure medications that act by blocking voltage-gated sodium channels, including carbamazepine, phenytoin, and valproic acid, had modest effects in our assay, even at elevated concentrations. It is currently unclear whether these differences in assay sensitivity are related to mechanisms of drug class, or the weak potency of the Nav blockers that we tested.

The primary screen, followed by secondary screen validation, demonstrated that this approach is robust and reproducible, particularly for detecting compounds with anti-convulsant properties. Given the added biological complexity of using live organisms, we found reproducibility to be strong, with ∼80% of the anti-convulsant compounds confirmed in the secondary screen. Therefore, this application is well-suited to efficiently and reliably screen large numbers of compounds to identify potent and efficacious compounds with anti-seizure properties. Previously, locomotion of zebrafish larvae has been commonly used as a surrogate measure of excitability to test anti-seizure drug effects (Baraban et al., 2013; Hoffman et al., 2016). Locomotor activity is a robust output but is an indirect readout of neural activity wherein compounds may impact motivation to swim or muscle physiology. Here, our use of a genetically encoded calcium sensor (expressed in neurons) provides a more direct measure of neural activity. In addition, our approach enables screening of a large number of compounds using a simple experimental readout that integrates drug effects throughout the assay. A drawback of using the plate-based approach for CaMPARI measurements is that the dynamic range of changes in the red:green ratio after convulsant treatment (∼2-fold) is lower than can be achieved with more deliberate manual confocal imaging of individual neurons labeled with CaMPARI, or other calcium sensors (e.g., GCaMP and GECO), patch-clamping a neuron directly, or locomotor activity (Baraban et al., 2005; Akerboom et al., 2012; Moeyaert et al., 2018; Zarowny et al., 2020; Gaudreau and Bui, 2025). Additionally, many of these assays provide a more detailed assessment of time-resolved effects of drugs, although this can add complexity to analysis and comparisons between drug effects.

Our tool can identify modulators of neural activity in the whole intact nervous system. Confirmed hits primarily targeted neurotransmission, as might be expected given the restricted expression of CaMPARI in neurons. Prominently affected signaling pathways included adrenergic, cholinergic, dopaminergic, serotonergic, GABAergic, and glutamatergic pathways. Not surprisingly, identified anti-convulsants included compounds that enhanced inhibitory GABAergic (e.g., ganaxolone) or suppressed excitatory glutamatergic pathways (e.g., (-)MK-801). The GABA-A receptor antagonist (+)-hydrastine was identified as pro-convulsant in our assay, aligning with the anti-convulsant effects of GABAergic enhancers. One of the identified anti-convulsants that is often not considered to target neurotransmission is the hormone progesterone. Progesterone is known to have antiseizure properties via its metabolite, allopregnanolone (Reddy et al., 2004; Miziak and Czuczwar, 2023). However, the exact target site is not well known, since traditional nuclear progesterone receptors would require longer to act than the treatment duration in our experiments. A recently identified cell-surface receptor, the membrane progesterone receptor γ (paqr5b), has been identified in zebrafish and may explain our observation, but needs future investigation (Mustary et al., 2024).

Our *in vivo drug* screening tool may be helpful in developing treatments for seizures in a variety of ways. Firstly, this assay can detect potent/efficacious compounds targeting hyperexcitability *in vivo*. Most drugs fail in animal studies and our innovation may provide a significant shortcut towards therapeutics for epilepsy. Secondly, as shown in Table 1 and Figure 6A, various drugs are already FDA-approved for conditions such as depression, arrhythmia, pain, and more. Therefore, some drugs may be considered for repurposing for treating seizures. Thirdly, several compounds may serve as templates for optimization. Notably, our screen identified several neuroactive steroids that could serve as templates for improvement, but are related to traditional anti-seizure drugs in that they function by activating GABA-A receptors (Sills and Rogawski, 2020). While targeting GABAergic systems for epilepsy remains a viable avenue, other targets should also be considered to provide novel options. Finally, as shown in supplementary figure 3, “Good Hits” refers to compounds that diminish effects of the 4-AP convulsant treatment, but have negligible effects under basal conditions. This may indicate strategies that target neuronal hyperexcitability during seizures while sparing activity under non-seizure conditions.

In summary, we have developed an *in vivo* drug screening assay that efficiently quantifies neural activity in thousands of individual larvae, and identifies compounds with anti-convulsant properties. Our CaMPARI assay will be useful for translational studies of conditions characterized by abnormal neuronal excitability, including epilepsy, pain, and other diseases involving abnormal neuronal firing. Our zebrafish CaMPARI assay will provide information on a drug’s performance in a physiological context. In the future, our tool can be combined with newly developed CRISPR-based technologies to efficiently generate zebrafish lines with altered disease genes (François et al., 2021; Parvez et al., 2021; Locubiche et al., 2024). Our CaMPARI-based *in vivo* screening approach may facilitate the identification of novel therapeutic targets for epilepsy while elucidating the mechanisms by which diverse signaling pathways modulate neural network dynamics and neuronal excitability.

## CONFLICT OF INTEREST DECLARATION

The authors have no conflicts to declare.

## DECLARATION OF TRANSPARENCY AND SCIENTIFIC RIGOUR

This Declaration acknowledges that this paper adheres to the principles for transparent reporting and scientific rigour of preclinical research as stated in the BJP guidelines for Design and Analysis, and Animal Experimentation, and as recommended by funding agencies, publishers and other organizations engaged with supporting research.

## ACKNOWLEDGEMENTS

This research was funded by a project grant from the Canadian Institutes of Health Research to Harley T. Kurata and W. Ted Allison (FRN 183619), and an NSERC Discovery Grant to W. Ted Allison. Ethan Smith was supported by an NSERC USRA.

## List of Abbreviations

4-AP: 4-aminopyridine
ANOVA: analysis of variance
CaMPARI: calcium-modulated photoactivatable ratiometric integrator
DMSO: dimethyl sulfoxide
dpf: days post-fertilization
GPCR: G-coupled protein-coupled receptor
LOPAC1280: library of pharmacologically active compounds 1280
Kv: voltage-gated potassium channel
KCNQ: potassium channel subfamily

---

**Supplementary figure 1.**
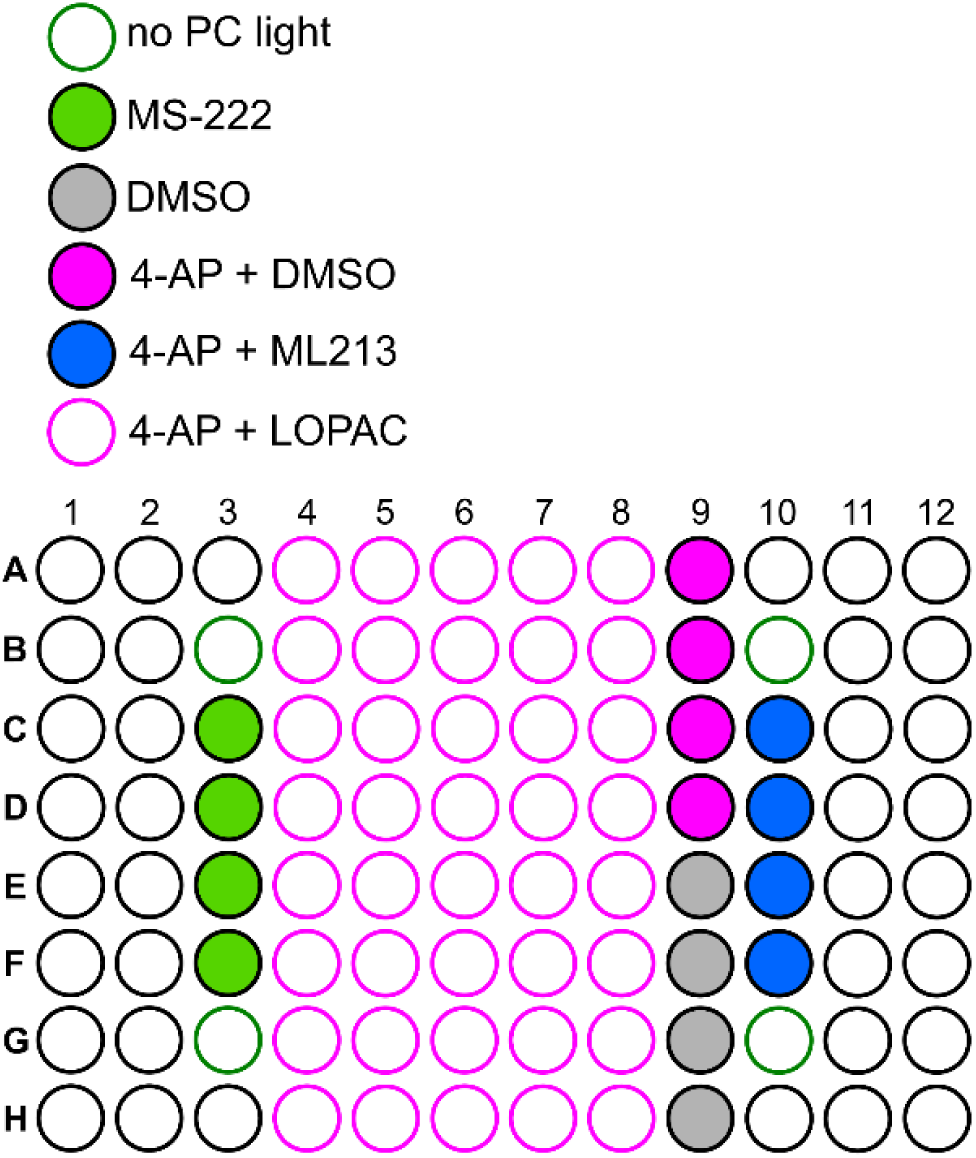
Representative 96-well “working” plate layout for zebrafish drug screen using CaMPARI. Zebrafish larvae (4 days post-fertilization) were placed in up to 64 wells of the 96-well plate and were exposed to the indicated conditions. The centre 40 wells included LOPAC test compounds. The various control treatments had up to four wells at the indicated conditions and locations. Note: Not all plates contained MS-222-treated wells.

**Supplementary figure 2.**
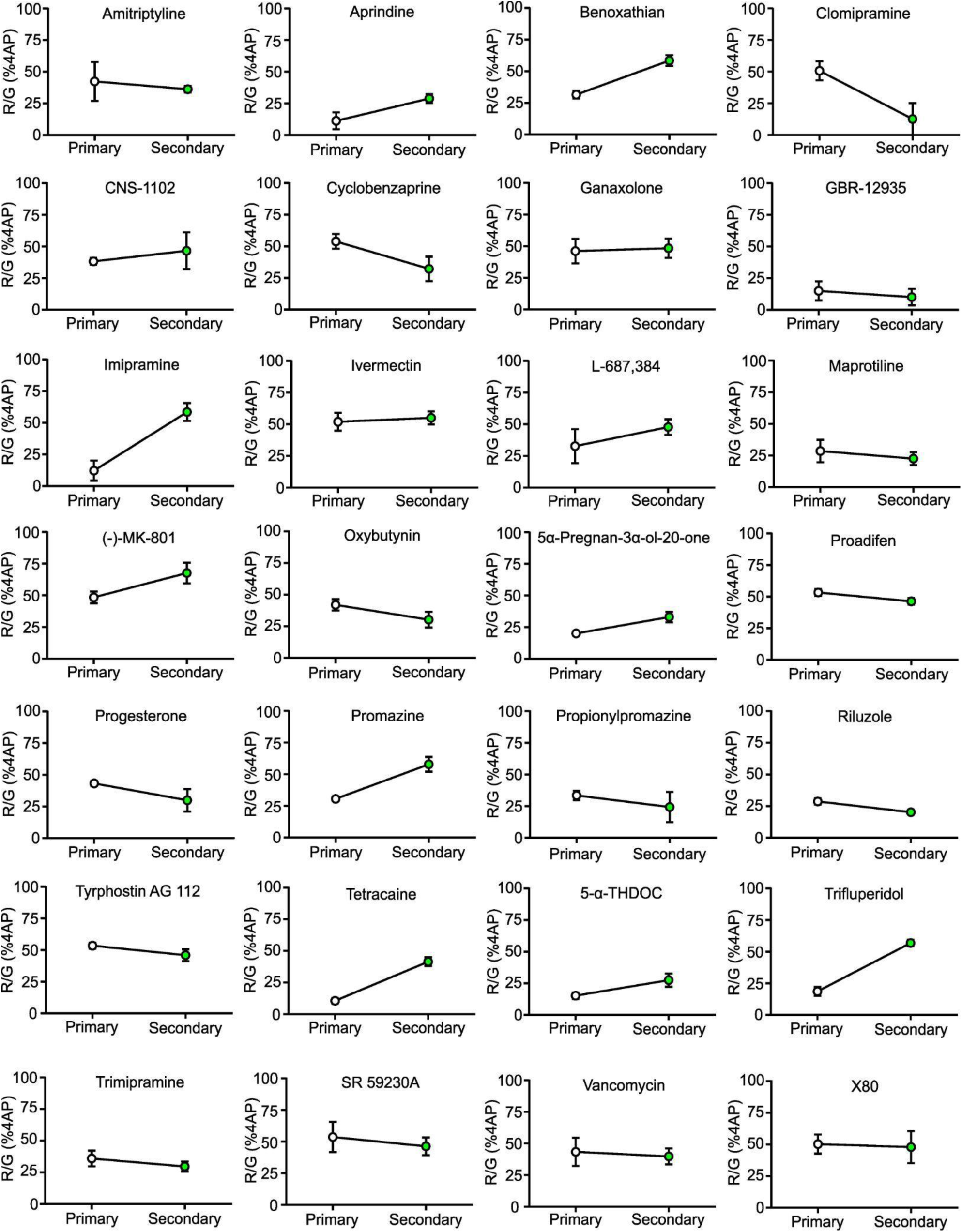
The reproducibility of anti-convulsants between primary and secondary screens. Zebrafish larvae (4 days post-fertilization) were treated with LOPAC1280 test compounds t (30 μM) for 30 min in the presence of 4-AP at 800 μM. R:G rations for all confirmed anti-convulsants in primary and secondary screens are presented to visualize the reproducibility of effects. Error bars indicate ± SEM.

**Supplementary figure 3.**
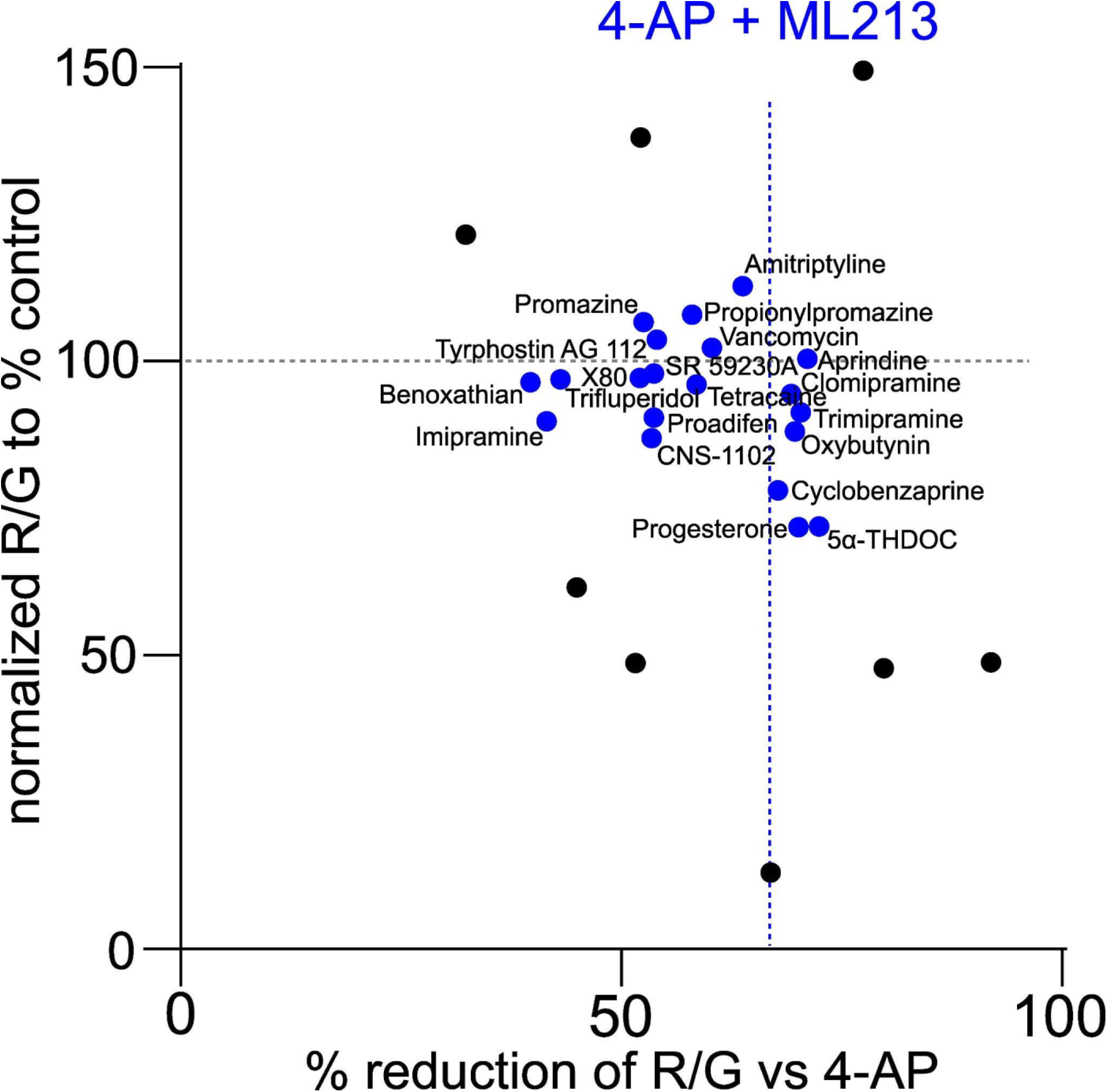
Identification of “Good Hits” as compounds with strong anti-convulsant properties but negligible impact under basal conditions. Compounds co-treated with 4-AP versus basal conditions in the absence of 4-AP. Most of the drugs identified are most effective in the 4-AP condition.

**Supplementary figure 4.**
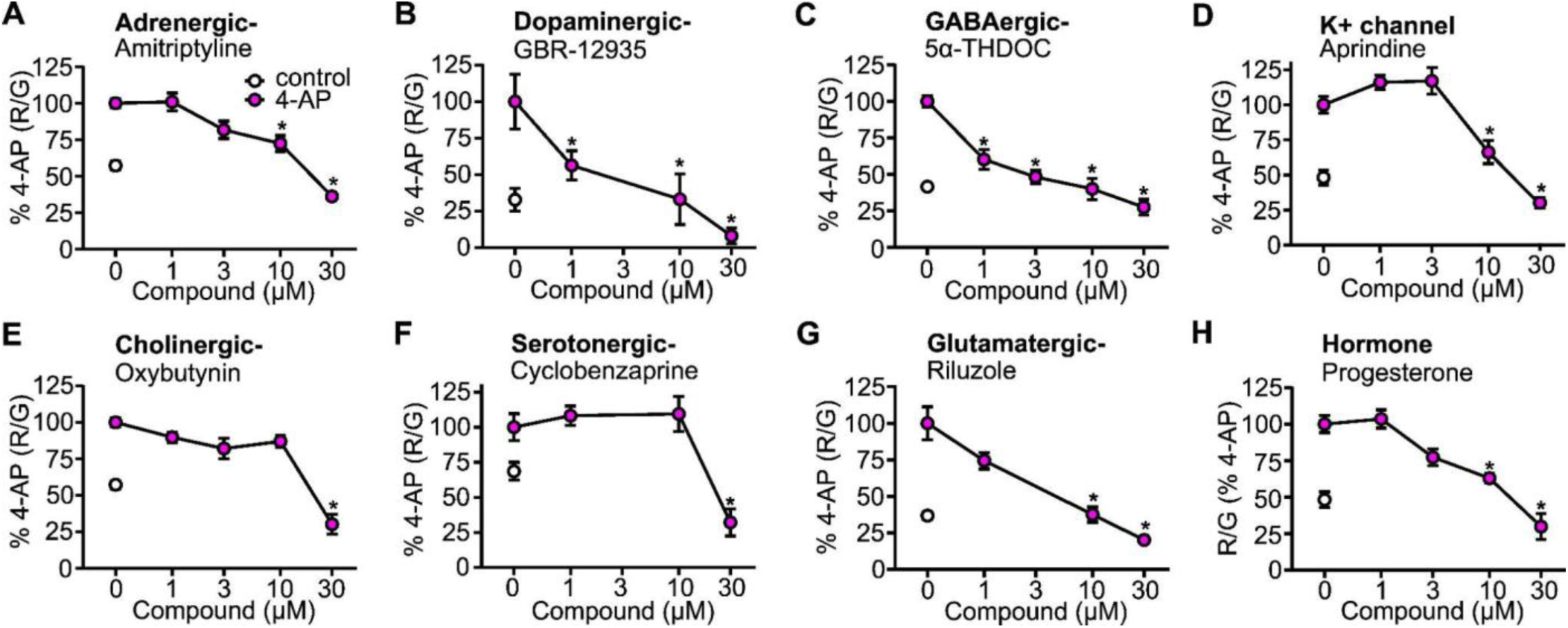
Anti-convulsant concentration-responses of confirmed hits. Indicated compounds were tested at various doses in the presence of 4-AP (800 μM) for 30 min. The R:Geratio was normalized to the 4-AP plus DMSO condition. ANOVA followed by Dunnett’s post-hoc test was used to determine statistical significance (P < 0.05). *, indicates statistical significance vs 4-AP alone. Biological replicates are at n=5-26.

## Notes

### Competing Interest Statement

The authors have declared no competing interest.

